# Ephaptic coupling and power fluctuations in depression

**DOI:** 10.1101/2025.11.15.688471

**Authors:** Dimitris A Pinotsis, Sankaraleengam Alagapan, Parisa Sarikhani, Tanya Nauvel, Christopher J Rozell, Helen S Mayberg

## Abstract

The initial therapeutic exposure to DBS during implantation surgery has reproducible acute behavioral effects that carry over without further stimulation. We analyzed LFP data from the first month following brief therapeutic intraoperative DBS. Data were recorded from the subcallosal cingulate cortex (SCC). During this month no further stimulation was applied. Recent studies have identified beta power fluctuations in LFP data as an acute putative depression biomarker of this exposure. However, a detailed description of neural dynamics underlying brain power fluctuations is missing. Here, we consider how these fluctuations are related to brain itinerancy, that is neural activity changes between stable and unstable states. We also provide a proof of principle study that these dynamics can be described using two new dynamical systems measures: instability frequency and relative wandering time. These capture interactions between neural activity and the mesoscale oscillatory electric fields generated by it. The two measures seem to split low vs. high HDRS scores within a small patient cohort. They are motivated by the cytoelectric coupling hypothesis, that suggests that efficient information processing results from mesoscale electric fields; and that the re-emergence of depression symptoms might result from altered electric fields. Whether the new measures reflect general mechanisms of rapid antidepressant action remains to be tested.

## Introduction

The brain’s oscillatory power is an acute putative biomarker in depression ^1^. Beta power fluctuations predict short term clinical effects ^2^. Early post-op theta cadence predicts antidepressant response five months later^3^. In brief, fluctuations in brain power seem to relate to depression state. However, a detailed description of neural dynamics underlying such fluctuations is missing. We here describe this dynamics using dynamical systems measures motivated by the cytoelectric coupling hypothesis^4^, that suggests that optimal information processing emerges from stable mesoscale electric fields, and that the recurrence of depressive symptoms may reflect a disruption of these fields.

DBS chronic treatment with ongoing stimulation can help maintain antidepressant effects for longer as a result of circuit stabilization^1^. Prior work has also shown that even brief intraoperative DBS delivered at the planned therapeutic target induces acute and reproducible symptom changes in core depression features^2^,^5-7^. In this setting, DBS functions not merely as a treatment but as a physiological probe— perturbing a pathological circuit away from its disease-associated state. What remains unclear, however, are the network dynamics that unfold following this short-term stimulation. Understanding how neural activity evolves in the weeks after brief intraoperative DBS may offer critical insights into the mechanisms by which acute modulation transitions to durable therapeutic benefit, and how short DBS perturbations can reorganize dysfunctional brain circuits to restore affective balance. To this end, we present a proof-of-principle study examining the network dynamics of the subcallosal cingulate cortex (SCC) during the first month following intraoperative therapeutic DBS^1^. During this period, no additional stimulation was applied. We considered SCC because it is a critical hub in depression, the target of DBS, and links to mood dysregulation, excessive rumination and treatment resistant depression^5–7^. Hyperactivity in the SCC correlates with depression severity, while reducing SCC activity through DBS or drugs is associated with significant mood improvement^8^.

Neural dynamics underlying SCC power fluctuations have been previously described using computational^9,10^ and machine learning^1^ models. Here, we used a dynamical systems model that describes the electric field generated by neural activity^11^. Based on this model, we then introduced two new dynamical systems–based measures—relative wandering time and instability frequency —to characterize fluctuations in neural activity and their interactions with mesoscale oscillatory electric fields. These metrics extend beyond traditional spectral measures, such as beta power, which have been proposed as acute biomarkers of antidepressant response. Relative wandering time is the relative frequency of electric field changes between stable and unstable states compared to neural activity, while instability frequency is the percentage of time that neural activity and electric fields are simultaneously unstable.

Our results suggest that these dynamical measures can distinguish between low and high HDRS (Hamilton Depression Rating Scale) changes within a small patient cohort over the month. Grounded in the cytoelectric coupling hypothesis^4^, which posits that efficient information processing depends on the integrity of mesoscale electric fields, our findings raise the possibility that the re-emergence of depressive symptoms may reflect disruptions in these fields and their effects on neural activity. Whether these dynamical signatures represent a general mechanism of rapid antidepressant action remains to be determined.

This paper comprises the following sections. In *Methods*, we present a mathematical model of ephaptic interactions and provide some tools needed to carry out dynamical systems analyses of brain power fluctuations. In *Results*, we characterize brain itinerancy, that is alternations between stable and unstable states over the month following surgery. We also consider ephaptic interactions underlying brain power fluctuations and discuss dynamical systems-based potential depression biomarkers. These results are further explained in the *Discussion*.

## Methods

### Participants and data

We reanalysed data from four subjects with treatment-resistant major depressive disorder published in ^1^. Ethics and experimental details are included in that earlier work. We report analysis of local field potentials (LFPs) during a period of four weeks after DBS treatment in the operating room. Bilateral electrode array leads were implanted in each participant, one in each subcallosal cingulate cortex (SCC) as determined from tractography previously described in^12^.

### Detrended Fluctuation Analysis

We analysed brain power fluctuations—a potential depression biomarker^1^ observed in our data. To characterize their dynamics, we looked at how they related to brain criticality. This is thought to balance order and randomness that might underlie mood regulation and cognitive flexibility in controls^13^. Brain power fluctuations have been linked to criticality in controls and excursions from it in depression and other patients^13–15^. Consistent with this view, markers of scale-free activity like long-range temporal correlations, are robust in wakeful resting EEG/MEG and degrade when the system shifts state (e.g., with sleep deprivation or anaesthesia) away from critical dynamics^16^,^17^.

To study critical dynamics and excursions from it, we used Detrended Fluctuation Analysis (DFA). This identifies and quantifies the presence of long-range correlations in LFP data ^18^. The primary goal of DFA is to detect a scaling exponent, often denoted as *α*, that characterizes correlation. DFA has been used in physiology, finance, geophysics, and many areas of science and engineering, where signals may exhibit correlations over long timescales^19,20^.

To find the scaling exponent, DFA uses the following steps: It begins with the calculation of the cumulative sum of the detrended time series (subtracting the mean from the original time series data). The integrated time series is divided into boxes of equal size, *n*, where *n* varies to observe the scaling over different timescales. This produces non-overlapping windows of equal length. In each window, a polynomial of a certain order is fitted to the data (typically linear, but higher orders can be used for more complex trends^18^). The fitted polynomial is then subtracted from the data to remove local trends. To sum, DFA starts with polynomial fitting and detrending. Then one calculates the root-mean-square (RMS) fluctuation: For each window size, the (RMS) term is calculated over all windows, providing a measure of how much the time series deviates from the local trends. Finally, the scaling behaviour is studied by plotting on a log-log scale against the size of the window. The slope of a line in this log-log plot is the scaling exponent, *α*, that characterizes the correlation properties of the time series.

### Ephaptic coupling in a neural ensemble

Above we discussed critical dynamics in brain power fluctuations. However, a detailed description of the evolution of these fluctuations is missing. To address this, in *Results* we introduce two dynamical systems measures. These are based on a combination of electric fields and Lyapunov exponents. The latter are discussed in the next section. Here we discuss electric fields. We consider a neural ensemble comprising pyramidal cells whose dendrites extend parallelly. (Supplementary Figure 1A). In this setting, branched dendrites are represented by cylindrical fibers. This symmetry aligns with the current dipole approximation widely used in human electrophysiology. Pyramidal neurons, aligned to produce an EF parallel to apical dendrites, receive synchronous input, generating dipole sources ^21,22^. They are combined into a unified extracellular space using the principle of superposition. Following up on earlier work ^11^ we used the *bidomain model* ^11^ to calculate the electric field (Supplementary Material and Supplementary Figure 1B). This model assumes that the electric potential discontinuity, generated by synaptic activity, gives rise to field sources. Despite their complex geometry, the parallel alignment of dendrites justifies the cylindrical fiber representation ^23^. This approach aligns with the current dipole approximation commonly used in human electrophysiology ^24^.

The model’s symmetry allows the extracellular field and potential to depend on just two spatial variables, *z* and *y*, instead of three. The *z* variable locates positions along the cylinder’s axis, while *y* represents a direction orthogonal to this axis. Because we here have recordings from only one electrode, this model can be further simplified to a classical model of an electromagnetic disc source (Supplementary Figure 1C). This is a reduction of the bidomain model considered above, where the axial length is negligible – corresponding to zero inverse conduction velocity, see ^25^. This is a common assumption that reduces neural field models to widely used neural mass models in Dynamic Causal Modeling (DCM) and elsewhere^26–28^, here used to simplify the bidomain model for the electric field. The electric field generated from a charged disc of circular radius is given by the following expression ^29^

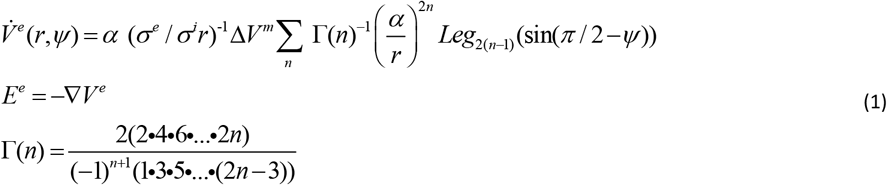

where *Leg*_*n*_ is the Legendre polynomial of order *n*. In *Results*, we use Equation (1) to calculate the electric field and using LFP data as a proxy for the transmembrane potential *V*^*m*11^. The disc source model approximates the electric field generated by a neural ensemble occupying a small patch in SCC by assuming that charge is uniformly distributed over a circular disc with radius *α*.

Having concluded the description of the electric field model that will be used in *Results*, we now consider an alternative expression to the bidomain model derived previously in^30^. This is the same as the bidomain model (Supplementary Material) with the addition of rate constants 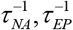. This will be clarified below. We did *not* use this alternative expression in *Results* because the addition of rate constants would bias our findings. On the other hand, using this alternative equation allows us to make a specific prediction about the stability of neural activity and electric fields —that cannot be obtained using Equation (1).

For simplicity, we assume that the LFP electrode is at a large distance compared to the size of the neural ensemble: the radius *a* of the fiber separating the intra- and extra-cellular spaces (grey cylinder in Supplementary Figure 1B) is very small compared to the vertical distance *y* to the location of the LFP electrode, *a* << *y*. This is shown by a squashed grey cylinder in Supplementary Figure 1D. The theory of electromagnetism ^29^ suggests that the temporal evolution of the electric potential to its resting value is given by

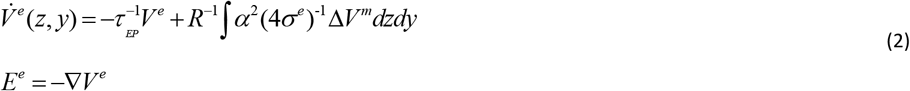

This equation furnishes the relaxation of the electric potential field at a rate 1/*τ*_*EP*_ within the bidomain model. In this model, the electric potential is generated by charges dispersed along an electrical fiber, parameterized by *z*, arising from the membrane current 1 / *r*_*i*_ Δ *V*^*m*^. The elementary source volume element is represented by *dzdy*, with *R* being the distance from the current source to the measurement point of *V* ^*e*^ in the extracellular space. The conductivity *σ* ^*e*^ of the extracellular space and the fiber radius are also included in the extracellular potential.

In ^30^, after expanding the integrand of Equation (2) in multipoles based on Legendre polynomials, the extracellular potential was found to obey the following equations

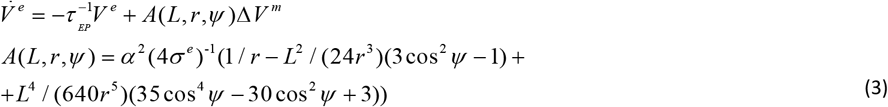

where *L* is the length of the neural fiber (occupied by the neural ensemble) and *r,Ψ* are called polar coordinates (Supplementary Figure 1C) an alternative to the usual Cartesian coordinates *x=rcosψ,y=rsinψ*. For the details of this derivation, the interested reader is referred to ^30^. To sum, Equation (3) describes how the extracellular potential unfolds in time before reaching a stable attractor as a result of neural ensemble activity, characterized by the membrane potential *V*^*m*^.

In^30^ it was also shown that the electric field Equation (3) can be coupled with an Equation describing neural activity. Considering a Fourier expansion of the transmembrane, *V*^*m*^, and extracellular potential, *V* ^*e*^ given by 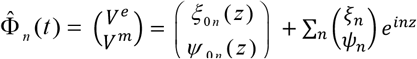 one can obtain an evolution equation that describes how how potential amplitudes unfold over time

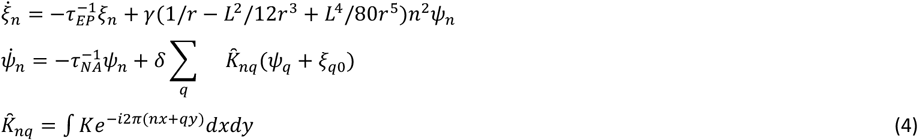

Above, we assumed the extracellular and membrane potentials are a superposition of harmonic waves *e*^*inz*^. Each wave described a potential with spatial periodicity. The amplitude coefficients (also known as *modes*), *ξ*_*n*_ and *ψ*_*n*_, characterise the amplitude of extracellular and membrane potentials. For mathematical convenience, we assumed that the point at which the extracellular potential is measured is vertical to the ensemble and at fixed distance. A similar result holds for any other angle.

Equations (4), describe ephaptic coupling, that is the interaction between electric field and its sources, spiking neurons. Note that a common assumption in bio-electromagnetism is that the electric field is quasi-static; the tissue reactance is assumed to be negligible and electromagnetic propagation effects can be ignored ^31^. This suggests that the time constants characterizing excursions from the stable state in Equations (4) satisfy the inequality 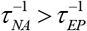.Then Equations (4) predict that the extracellular potential modes *ξ*_*n*_ are more stable than the faster relaxing neural activity modes *ψ*_*n*_ ^32^.

All in all, we hypothesized that brain power fluctuations might correspond to changes in the brain’s electrical activity at both the neural and mesoscale electric field level. We discussed the coupling between these two levels (Equation 4) and considered an equation for the electric field (Equation 1). Our mathematical analyses suggested that neural activity is more unstable than the field activity. In *Results*, we will test this prediction and also use the electric field equation to predict electric field responses and ephaptic coupling (i.e. neural to mesoscale electric field coupling) that might underlie power fluctuations.

### Circular causality and electric field—neural activity interactions

Above, we considered ephaptic coupling. In ^30^ it was shown that ephaptic coupling is an application of the more general slaving principle in Synergetics^30,32^. Following this principle, fast relaxing quantities, like the membrane potential modes *Ψ* _*n*_, depend on slowly varying quantities, like the extracellular potential Coefficients *ξ* _*n*_, appearing in Equation (4) above. The first mode of neural activity *ψ*_*1*_ is enslaved by the mode *ξ*_*1*_ of the extracellular potential (and similarly for higher modes). Indeed, from the second line of Equations (4), it follows that

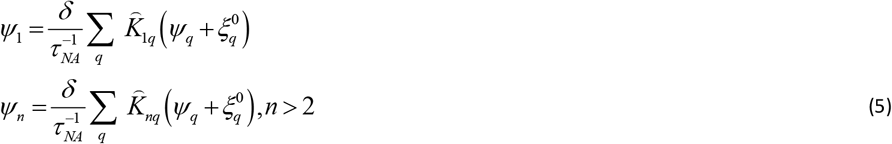

where *q* indexes the modes, 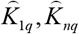 are defined in Equation (4) and 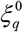 is the boundary value of 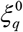 on the grey cylinder of Supplementary Figure 1B. The slaving of the membrane potential mode *ψ*_*1*_ by the extracellular potential modes *ξ*_*n*_ can also be written as

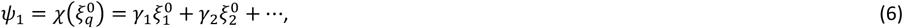

where the coefficients *γ*_*1*_, *γ*_*2*_ etc are given in terms of the constants 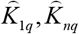, *δ*, and *τ* _*NA*_ that can be determined using algebra. Equation (6) is readily obtained from Equations (5) after substituting the second line in the first. Having obtained an expression for the transmembrane potential mode in terms of the extracellular potential modes we can now close the loop by using Equation (6) together with the first of Equations (4) (for *n=*1) that expresses the reverse, i.e. the extracellular mode in terms of the transmembrane. This reads as follows

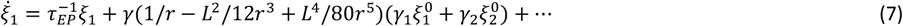

Because of the terms involving 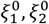 etc, Equation (7) suggests that the evolution (change) of the extracellular potential depends on the value of the *near field*, 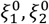 etc,that is, the electric field very close to the ensemble (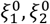 are the boundary values of the modes 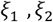 etc). Electric field changes on the boundary of the neural ensemble 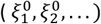 as a result of some external input to the cortical patch or of DBS. Then, the extracellular potential mode *ξ*_*1*_ predicted by Equation (7) will change. At the same time, the source, that is the transmembrane potential mode *ψ*_*1*_, will change simultaneously as predicted by Equation (6). To sum, Equations (6) and (7) predict that perturbing the electric field close to the cortical patch occupied by a neural ensemble changes the extracellular potential *and* its source (the neural activity that generates it) – which is circular causality.

### Lyapunov exponents

The new measures that will be introduced in *Results*, to describe SCC power fluctuations are combinations of Lyapunov exponents. Lyapunov exponents quantify the rate at which brain dynamics change over time. We calculated the largest Lyapunov exponent from the available LFP data and electric fields (computed with the model of the previous section). We started with time series and followed^33^ to find the largest Lyapunov exponent. This describes the system’s robustness, i.e. sensitivity to perturbations. Mathematically, it quantifies the average exponential rate of separation between nearby trajectories in the phase space. Intuitively, if the largest Lyapunov exponent is positive, small differences in initial conditions will grow exponentially over time, while if it is negative the system is stable to small perturbations and will return to a steady state or periodic orbit after a disturbance. Zero Lyapunov exponents suggest that small differences in initial conditions neither grow nor shrink over time. This can indicate neutral stability, where the system may have regular, periodic behavior. To obtain the Lyapunov exponents, we used the following steps:

For a time series of brain data *x*(*t*_*i*_),*i* = 1, 2,… the phase space was reconstructed following Taken’s theorem ^34^ using delay coordinates *[x(t),x(t+τ),x(t+2τ),…,x(t+(m−1)τ)]*, where *τ* is called delay and was chosen by minimizing mutual information between delay coordinates and the original time series, so that information is preserved. The parameter *m* is called embedding dimension and was chosen using the method of false nearest neighbours^35^, which suggests that if two points are close in a space of dimension *m*, but not do not stay close in a space of one more dimension *m+1*, they are false neighbours. The right embedding dimension was then found when the percentage of false neighbours was very small. After the phase space had been reconstructed, the largest Lyapunov exponent was obtained by evolving the distance between two nearest neighbours and tracking how much they diverge over time.

## Results

### Deviations from criticality follow HDRS scores

We analysed brain power fluctuations in LFP data recorded from SCC that have been shown to be a putative depression biomarker^1^. We characterized their dynamics using measures of brain criticality. Brain power fluctuations have been linked to brain criticality in controls and excursions from it in patients. Criticality suggests that the brain operates near critical points. These balance order and randomness^13^. This, in turn, is thought to optimize information transmission^36^, cognitive flexibility^37^ and metabolic consumption^38^. Criticality is thought to enable efficient information processing^16^, and help maintain mood stability^17^. Deviations from critical dynamics have been observed in diseases and disorders including depression^13–15^. Such deviations are manifest as e.g. rigidity, slowed cognitive processing, and mood swings— common symptoms of depression. This is accompanied by pronounced changes in brain dynamics. Depressed patients show hyperactivity in the default mode network^39^ and dysregulation in the salience network^40^.

We asked if critical dynamics and deviations from it could be found in LFPs recorded in depression patients during the four weeks following the OR treatment with DBS^1^. In the OR, a 10 minute session of bilateral optimized high frequency stimulation was administered. The subsequent four weeks of recordings were performed with the stimulation turned off. We analysed longitudinally collected data from the left and right subcallosal cingulate cortex (SCC) from four depression patients. These were previously studied in^1^. Before surgery, all subjects had Treatment Resistant Depression (TRD) and a Hamilton score (HDRS) above 20. After surgery and exposure to 10 minutes of therapeutic stimulation at the optimized target in the OR, HDRS scores dropped significantly. It is at this new state that the recordings we consider here started. These four subjects were chosen due to the differences in their HDRS scores across the four-week analysis period and to guarantee coverage of the full range of possible scores: Subject P001 had a low HDRS score that corresponded to mild depression (HDRS=4—8). P003 showed the largest increase in HDRS, moving from the mild to moderate range (9–16). Meanwhile, P004 and P002 had consistently higher scores, falling within the moderate-to-high or very high ranges (13–19 and 15–21). Despite its size, this subject set spanned all HDRS ranges within the brief observation period of this proof-of-principle study. In future studies, we will analyze a longer time window and a larger cohort.

We chose SCC because it is known that depression is associated with hyperactivity in SCC and a limbic-frontal network ^10,41^; SSC is also the target of neuromodulation^42^and has been associated with mood dysregulation and treatment resistant depression^5–7^. In all our analyses in this and later sections, we used data recorded from midnight till morning hours to reduce circadian variations. We used Detrended Fluctuation Analysis (DFA; *Methods*) to quantify critical brain dynamics^43,44^. DFA characterizes brain dynamics in terms of a scaling exponent, often denoted as *α*. This is the power of correlations *f* ∼ *t*^*α*^. It represents how fluctuations in a time series increase with time. For *α*=0.5, the time series corresponds to white noise. For 1>*α*>0.5 the time series corresponds to critical dynamics. It includes long-range temporal correlations (LRTC).

The scaling exponent, *α*, has been used as a marker for neurological diseases and disorders ^45,46^. Studies have reported differences in mean scaling exponent values between controls and depression patients. Often comparative results between groups are presented and scaling exponents are used to describe between group differences. Both higher (i.e. *a*>1) and lower (*a*<0.5) scaling exponents have been found in EEG data from depressed individuals ^47,48^. *α<* 0.5 corresponds to anti-correlations ^15^. These can be thought of a mechanism imposing rigidity in the system producing the time series, leading to fewer fluctuations than chance. For *α*>1 the process is thought to be non stationary—this has also been observed in EEG recordings from depression patients^48^. To sum, for 1>*α*>0.5 the brain shows critical dynamics. Values around 0.6-0.8 are often associated with controls^47^. Outside this range, the brain deviates from criticality. This has been observed in patient cohorts ^47,48^.

Scaling exponents are often averaged over the duration of a day to smoothen out intra day and circadian variations and artefacts^48,49^. Within day variability can be around 30% or more^48^. Calculation of the exponent is also sensitive to preprocessing and filtering choices^50^. To mitigate variability average measures are used. Below, we asked whether the median scaling exponents in our cohort deviated from criticality for each subject and every day. The results of our analyses are shown in Figure 1.

**Figure 1.**
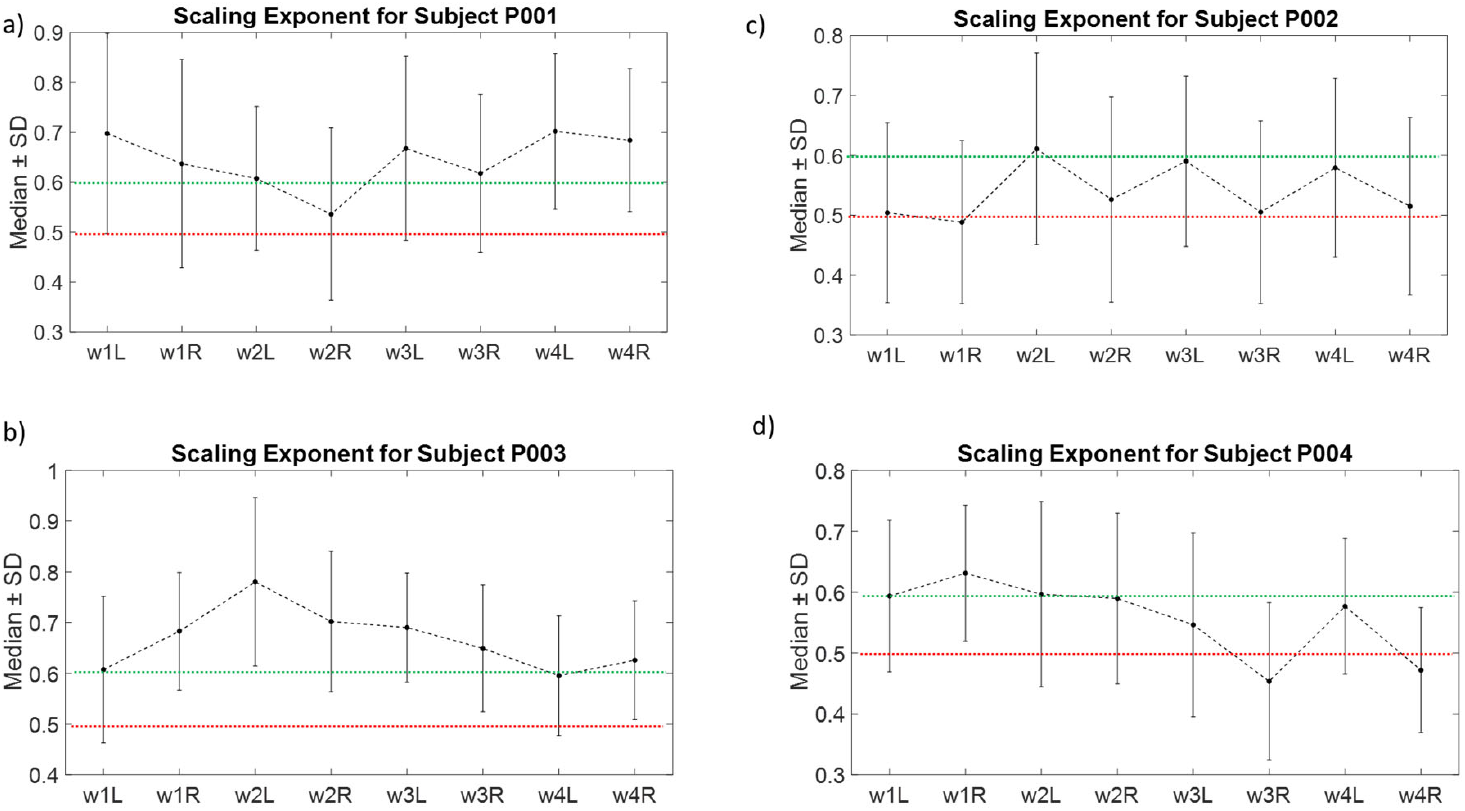
Median and Standard Deviation of Scaling Exponents Over Sample Recordings for Four Subjects. Panels a)–d) display the median scaling exponents, α, and their standard deviation, calculated from sample recordings from left and right hemispheres during the four weeks following OR. Each panel represents data from a different subject, illustrating variations in the median scaling exponent,α, on the y-axis. Each data point corresponds to an individual α estimate averaged across a week based on daily recordings. The horizontal red dashed line corresponds to the critical value 0.5 that defines white noise, while the green dashed lines depict the values 0.6 and 0.8. This is the range often associated with controls. HDRS scores are also shown on the horizontal axes alongside the week and hemisphere that corresponds to each estimated value of the scaling exponent.

Panels a)-d) show the median scaling exponents for Subjects P001,P003, P002 and P004. They show average values for each week during the first month post OR for the left and right hemispheres. Interestingly, HDRS scores agree with scaling exponent medians. Recall that Subject P001 had a low HDRS score (4—8). The subject had very little return of symptoms over the period considered here. This was confirmed by the scaling exponent shown in Figure 1a). It showed the smallest deviations from criticality. The median was consistently above 0.6 (except in week 2 for the right hemisphere), which is a value similar to that often observed in controls. This agrees with the low HDRS score.

Recall also that Subject P003 had moderate depression (HDRS=9—16). Figure 1b) shows the corresponding median scaling exponents. They range between 0.6—0.8 (the range between the two green dashed lines often associated with controls). Last, as mentioned above Subjects P002 and P004 had the highest HDRS scores (15—21 and 13—19). Here the median scaling exponents fluctuated close to the critical value *α*=0.5, see Figures 1c) and 1d). They ranged between correlated and anti-correlated states, i.e. above and below the red dashed line. Similar values for *α* (around 0.5) have been linked to psychomotor retardation and dopaminergic deficiency in depression ^51^,^47,48^.

To sum up, we found fluctuations in the median scaling exponent during the first four weeks after OR. These were in accord with HDRS scores. Higher HDRS scores were associated with larger deviations of the scaling exponent from criticality ^14,15^. This is also reminiscent of correlations between HDRS scores and median scaling exponents found elsewhere^48^.

### Lyapunov exponents measures reflect HDRS scores

Above, we found deviations from critical brain dynamics in depression subjects. These deviations were quantified in terms of scaling exponents. We then asked if brain power fluctuations and excursions from criticality considered above might be the result of changes in the mesoscale oscillatory electric fields generated by neuronal spiking. The idea is that electric fields turn around and affect neuronal spiking and possibly electrochemical processes in the brain’s cytoskeleton. This is motivated by a recent hypothesis known as cytoelectric coupling^4^. This suggests that i) efficient information processing results from top down mesoscale field influences and ii) that cognitive deficits of the sort observed in depression might result from altered electric fields and variations in brain power. To test this hypothesis, we computed the Lyapunov exponents of neural and electric field responses. The latter were obtained using an electric field model described in *Methods*.

Lyapunov exponents depict the rate at which neural activity converges or diverges from an attractor state. The existence of such states (attractors) is a common assumption in cognitive neuroscience ^52,53^. They have been used in memory ^54^, perception ^55^ and decision making ^56^ and underlie oscillatory dynamics ^30,57^. Following that earlier work, we also assumed attractor dynamics in our resting state data. We analysed exactly the same data as in the previous section: local field potentials (LFPs) from SCC in depression patients. These were recorded during early morning hours over the four weeks following OR treatment^1^. For illustration purposes, Figure 3 shows the results of the first week. The quantitative results below (Figure 4) were based on data recorded from the four weeks month post OR (similarly to all analyses considered here).

For each subject, we computed two sets of Lyapunov exponents based on two different time series. The first used direct recordings of neural activity obtained with LFP; the second used predictions from our electric field model (disc source) that resulted from this neural activity (*Methods*). We thus obtained two sets of Lyapunov exponents: one corresponding to neural activity and another corresponding to electric fields. Motivated by mathematical arguments discussed in *Methods*, we predicted that positive exponents for neural activity will be larger than those for the electric field. This suggests that neural activity is more unstable than electric fields in accord with earlier results ^11,30^.

Our prediction was confirmed. This can be seen by going through Figures 2b)d)f) and h). These Figures show the Lyapunov exponents for patients P001,P002,P003 and P004 corresponding to neural activity and electric fields. Lyapunov exponents based on neural activity are shown in blue bars, while the corresponding exponents based on electric fields are shown in orange bars. Positive exponents are shown with bars facing upwards, while negative exponents are shown with bars facing downwards. Sample recordings from the first week after surgery are shown on the horizontal axis. For most positive exponents, blue bars are larger than orange, that is the Lyapunov exponents corresponding to neural activity were larger than the exponents corresponding to the electric field. Thus, neural activity is more unstable than fields. For illustration purposes (to avoid cluttered plots) in Figure 2 below we show the Lyapunov exponents over the first week only.

**Figure 2.**
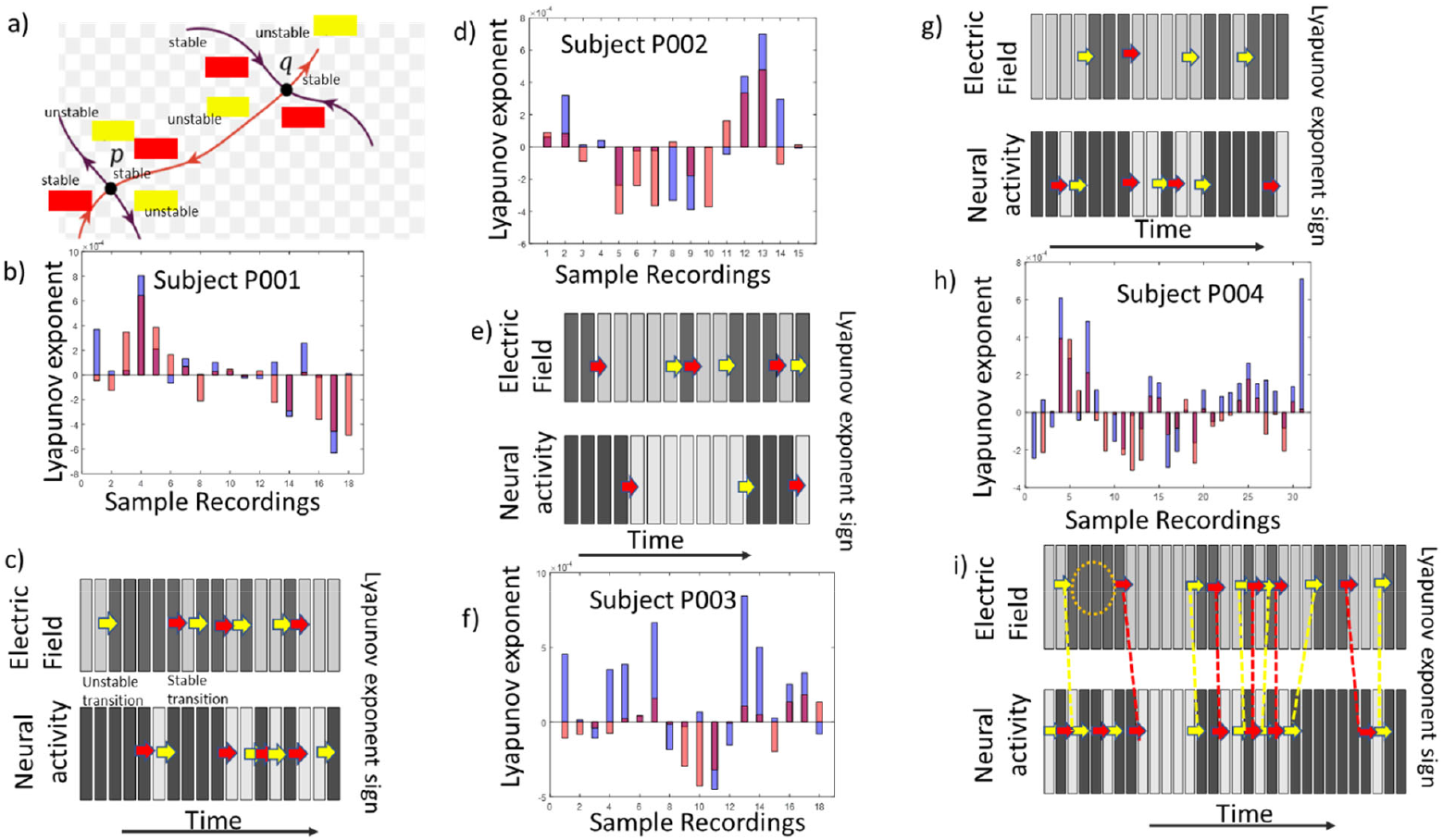
Lyapunov Exponents. Panel a) shows Itinerant dynamics in phase space. The points p and q are attractors (fixed points) and arrows depict stable and unstable directions. Fixed points are connected via heteroclinic orbits that illustrate itinerant dynamics, i.e. transition pathways between equilibria. The obits showcase how the system evolves, either stabilizing, diverging, or transitioning between states. Panels b),,f) and h) show the Lyapunov exponents over the course of the first week after OR. Blue bars correspond to Lyapunov exponents based on neural activity while red bars to exponents based on electric field. The horizontal axis includes the number of samples (2-3 / day for the first 7 days after OR). Panels c), g) and i) show the same data as panels b), d),f) and h). Positive exponents in panels b), d),f) and h) are shown with dark grey bars in panels c), e) g) and i). Negative values with light grey bars. The additional information in panels c), e) g) and i) is shown using red and yellow arrows. These depict stable and unstable transitions, i.e. changes in the Lyapunov exponent sign. See text for details.

Figures 2c)e)g) and i) contain the same information as Figures 2b)d)f) and h). They depict the signs of Lyapunov exponents separately for neural activity and electric fields (as opposed to being overlaid in the earlier plots). Exponents for electric fields are in the top panels and for neural activity in the bottom panels. Each bar is either light or dark grey. Light grey corresponds to a negative sign (stable dynamics; see *Methods*), while dark bars to a positive sign (unstable dynamics). Positive and negative exponents alternate over time. Alternating signs for Lyapunov exponents is a hallmark of itinerant dynamics, that is dynamics that move between attractors (stable states). This is what also happens here. Yellow arrows denote unstable transitions (from negative to positive exponents, diverging from an attractor), while red arrows stable transitions (from positive to negative, converging to an attractor).

To sum, alternating signs of Lyapunov exponents suggest that brain activity moves between attractors. This sort of dynamics is called *itinerant dynamics* ^58^. This result is also confirmed in the next section. To understand itinerant dynamics, consider the following analogy: a golf ball (representing a brain circuit) is rolling across a hilly golf course with shallow dips and slopes (attractors), see Figure 2a). The ball lingers in these dips, almost settling but not fully trapped, as the dips aren’t deep enough to hold it indefinitely. There’s always a slight slope that eventually pulls the ball out, defining a path known as a *heteroclinic channel*, with the dips representing *saddle points* (see *Methods* and ^59^). The dip centres are denoted as “*p*” and “*q*” in Figure 2a). In some directions (noted as “unstable” in the figure, also shown with a yellow rectangle) the ball is pulled out. Note this corresponds to a *positive* Lyapunov exponent. In other directions, the slope is steep and the ball cannot escape (noted as “stable” in the figure, also shown with a red rectangle). This corresponds to a *negative* Lyapunov exponent. Suddenly, the ball might roll up and over the ridge of the valley and heads toward another part of the golf course, driven by chaotic forces (like unpredictable gusts of wind or varying slopes), bouncing around. Each time the ball moves, it follows a different path, never exactly repeating the same motion, and never following a predictable path. Similarly, cortical dynamics in depression patients might wander between different attractors. This is chaotic itinerancy ^60,61^.

Based on the above calculation of Lyapunov exponents for neural activity and electric fields we now consider two measures that can describe altered ephaptic interactions in the four weeks following OR. First, relative wandering time. The frequency of sign changes (i.e. that the total number of red and yellow arrows) in each plot of Figure 2 provides an estimate of the time that the system wanders between attractors. We call this *wandering time*. Then, the *relative* wandering time between neural activity and electric fields (i.e. the total number of arrows in the top and bottom panels in Figures 2c)e)g) and i) describes the relative flipping count: the number of times the electric field alternates between moving away and toward a stable state after subtracting the corresponding number for neural activity. We computed the relative wandering time for all subjects and both hemispheres. This is shown using green lines in Figure 3 (*“DWandering Time”*). Solid lines indicated differences from the left hemisphere and dashed lines from the right hemisphere. Each plot consists of four points, one for each week. For convenience, at the bottom of the plots we included the HDRS scores for each week.

For subject P001, with mild depression the relative wandering time in the left hemisphere was always negative (ranges between -2% and –15%, cf. Figure 3a). The minus sign here and elsewhere suggests that the wandering time of electric fields was smaller than neural activity. Similarly, the wandering time for subject P003 with moderate depression scores was also negative in the left hemisphere (Figure 3c). Neural activity wandered more. In contrast for subjects P002 and P004 with severe depression, the relative wandering time was positive in the left hemisphere across almost all weeks (except one), see Figures 3e and 3g. Neural activity wandered less. To sum, low HDRS scores seemed to correspond to negative relative wandering time in the left hemisphere, while high HDRS scores to positive values of the same measure. Interestingly these values were obtained based on LFPS from the left hemisphere. We come back to this point below and in the Discussion.

**Figure 3.**
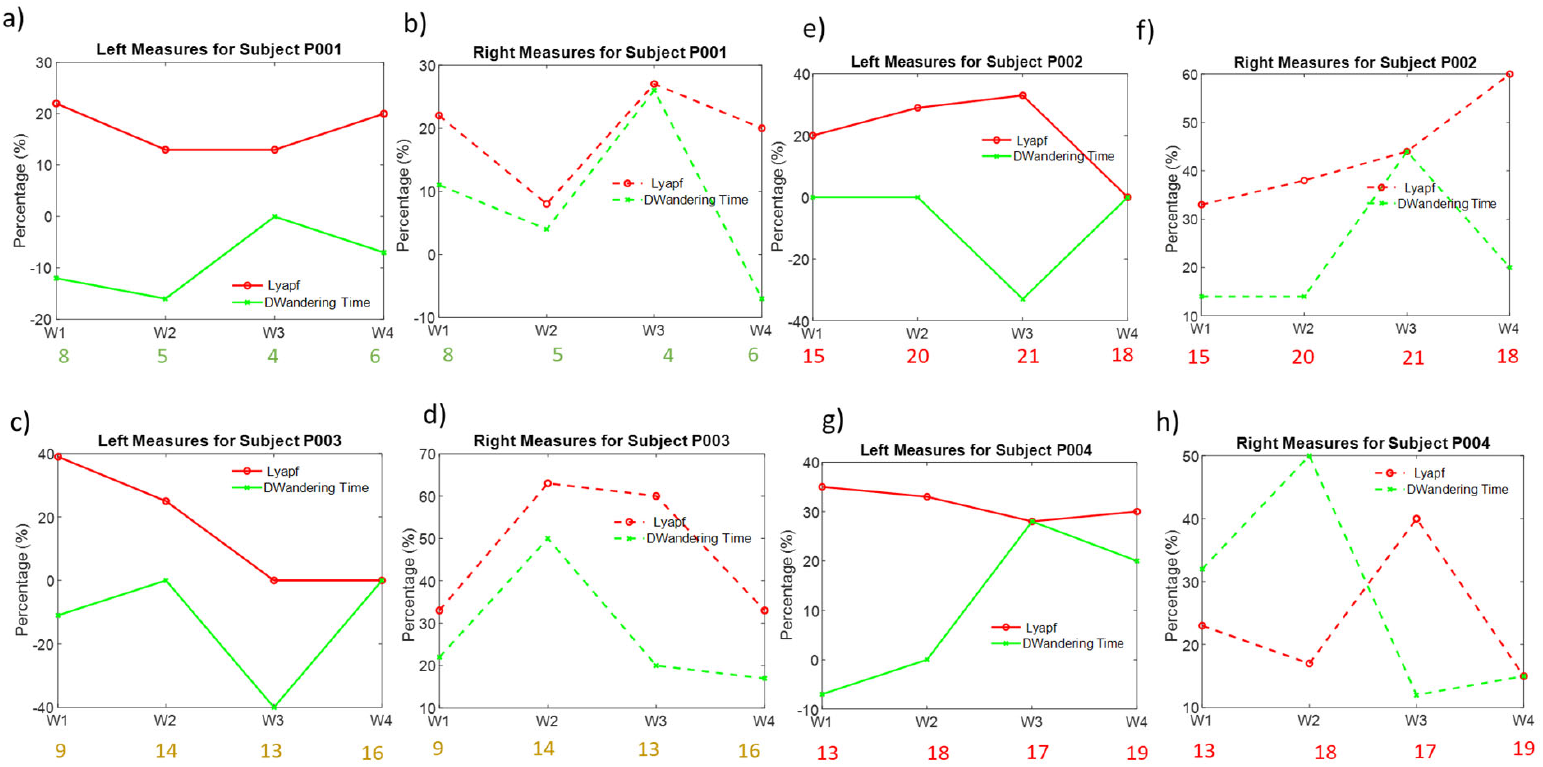
Relative Wandering Time and Instability Frequency. Panels (a–h) display data from four subjects, during the month following OR, arranged in left/right hemisphere pairs. For each subject, the left panel corresponds to the left hemisphere and the right panel corresponds to the right hemisphere: (a,b) Subject P001, (c,d) Subject P003, (e,f) Subject P002, and (g,h) Subject P004. Within each panel, instability frequency is shown in red and relative wandering time in green (see text for details). HDRS scores are shown at the bottom for the corresponding weeks.

Besides relative wandering time, we considered a second instability measure based on Lyapunov exponents. Instability frequency is the percentage of time that Lyapunov exponents for neural and electric fields were simultaneously positive. During these epochs, the brain moved away from the stable state in both the neural activity and electric field phase space. This measure is also shown using red lines in Figure 3 (*“Lyapf”*).

For Subject P001, the instability frequency in the left hemisphere ranged between 10-20% (red line in Figure 3a) Interestingly, for subject P003, that also had relatively low HDRS scores, the instability frequency in the left hemisphere dropped to zero in weeks 3 and 4 (red line in Figure 3c). Thus instability frequency between 0-20% in the left hemisphere seemed to correspond to low HDRS scores (with the exception of the first sample in subject P003). On the other hand, higher HDRS scores in subjects P002 and P004 seem to be associated with instability frequency above 20% in the left hemisphere (red lines in Figures 3e and 3g). To sum, it seemed that lower HDRS scores were associated with lower instability frequency, below 20%, while higher HDRS scores signified an instability frequency above 20%. Again, values in the left hemisphere seemed to suggest a split of the subjects based on high vs low depression score, similarly to relative wandering time above.

All in all, we characterised brain dynamics in depression patients using Lyapunov exponent measures. These seem to follow HDRS scores. First, relative wandering time, that is, the percentage of time that electric fields flip between stable and unstable states in relation to the percentage that neural activity flips. Here, mild and moderate depression was associated with negative values, while zero and positive values were observed in subjects with severe depression. Second, instability frequency, that is the time percentage that electric fields and neural activity were simultaneously unstable. Higher values (above 20%) seemed to be linked to severe depression, while lower values (around 20% or less) were observed in subjects with mild or moderate depression. Note that our taxonomy used values of both measures in the left hemisphere only. These seemed to be linked to HDRS scores. This is not an isolated choice. Many quantitative measures associated with depression severity seem to relate to brain activity in the left hemisphere. These are summarised in the Discussion section below.

### Lyapunov exponents correlate with brain power

Above we considered itinerant dynamics in depression patient data during the month after OR. Neural activity and electric fields seem to move between quasi-stable attractor states. We characterized itinerant dynamics using Lyapunov exponents. Earlier work considered in ^1^ found that fluctuations in oscillatory power in the same brain dynamics. Taken together, these results suggest that Lyapunov exponents might be linked to brain power changes. This is what we looked for in our final analyses. We asked if Lyapunov exponents correlated with oscillatory power obtained from LFP recordings. If so, this implies that Lyapunov exponents and brain itinerancy are linked to brain power fluctuations.

We computed correlations for all subjects for Lyapunov exponents obtained using the LFP data and found they were significant. These are shown in Figures 4a)-d). Lyapunov exponents based on neural activity were depicted on the horizontal axis. The vertical axis shows broadband power (1-70Hz in Figures 4a)-c) and beta power in Figure 4d). A multiple linear regression was performed to predict power from Lyapunov exponent while controlling for recording week. The model revealed a significant positive effect of Lyapunov exponent on power (β = 8.49×10^−7^, *p* = 1.2×10^−20^ for Subject P001, see Figure 4a)). The effect of week was negative and did not reach significance (β = –3.95×10^−11^, *p* = 0.068 for Subject P001). Similarly, for Subject P002 regression analysis revealed a strong positive association between *X* and power (β = 7.26 × 10^−6^, *t* = 11.25, *p* = 2.8 × 10^−19^), while the effect of week was not significant (β = 2.82 × 10^−10^, *p* = 0.70), see Figure 4b). For Subject P003 (Figure 4c) power increased significantly with Lyapunov (β = 2.75 × 10^−7^, *p* = 4.0 × 10^−6^), while week had no significant effect (*p* = 0.61). Last, for Subject P004 there was no significant dependence of overall power on Lyapunov exponents (*p* =.14). However, beta power increased as Lyapunov exponents increased (Figure 4d; β = 2.58 × 10^− 8^, *p* = 5.04 × 10^− 9^). Interestingly, the effect of week was also significant, β = 2.52 × 10^−11^, *p* = 3.21 × 10^− 7^.

**Figure 4.**
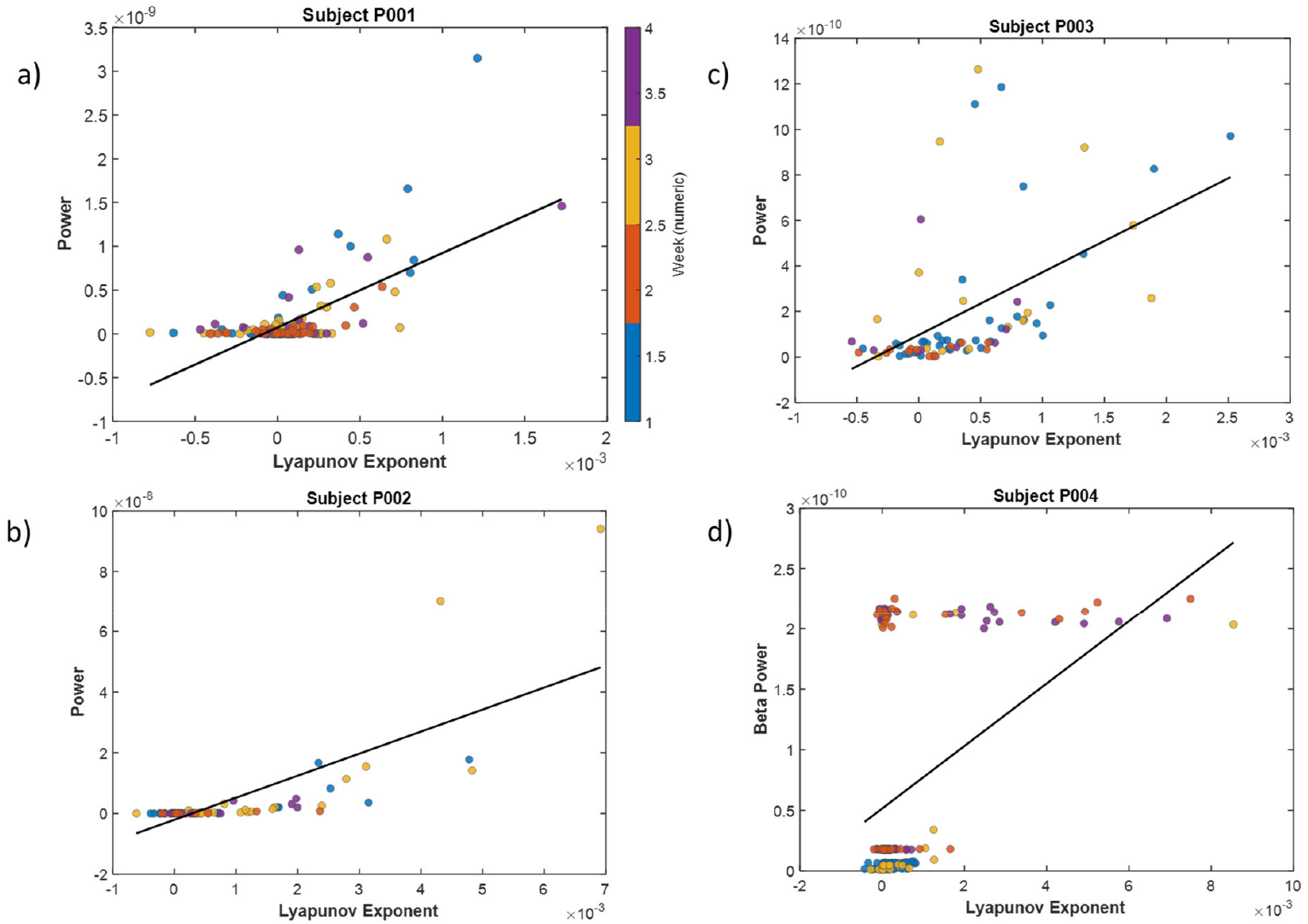
Brain Power Fluctuations and Itinerant Dynamics. Regression analysis showed a strong positive association between Lyapunov exponent and power, see text for details.

Overall, our results indicate that greater Lyapunov exponents are associated with increased power. For most subjects, results were independent week with the exception of P004 that showed a significant effect of Lyapunov exponents on beta power specifically. This analysis suggests a link between broadband power fluctuations and Lyapunov exponents. It confirms the result in the previous section that fluctuations in oscillatory power in our data are linked to itinerant brain dynamics.

## Discussion

We considered fluctuations of brain power during the first four weeks after brief intraoperative DBS in the operating room. These were observed in local field potential (LFP) electrode recordings from the subcallosal cingulate cortex (SCC) – and assumed that LFP recordings reflected recurrent SCC activity and afferent inputs from thalamus or limbic structures ^62^. We asked if the observed fluctuations corresponded to itinerant dynamics, where brain activity moves between different attractor states. We found that Lyapunov exponents based on recorded brain activity correlated with SCC oscillatory power. This suggests that power fluctuations might reflect itinerant dynamics as the brain wanders between different attractors.

We also asked if fluctuations corresponded to changes in the brain’s electrical activity. This was motivated by cytoelectric coupling^4^, a hypothesis that suggests that persistent symptoms in depression might correspond to altered mesoscale electric fields. This hypothesis also suggests that electric fields organize neurons to perform efficient information processing and considers ephaptic influences as necessary for coordinating neural activity. To describe top down field influences, we used a model that predicts electric fields based on LFP activity. This allowed us to characterize changes in mesoscale electrical activity as a result of the brain’s endogenous activity or carry over effects of intra operative DBS or both. We used the theory of electromagnetism in physics to predict the brain’s near field, i.e. the electric field very close to an SCC patch—and obtained a second time series on top of the LFP responses characterizing brain activity. Then, we described interactions between aggregate electric fields and neural activity.

To this end, we introduced two new measures that characterize ephaptic interactions. These included combinations of Lyapunov exponents. First, relative wandering time. This is the relative frequency that the electric field alternates between stable and unstable states with respect to neural activity. Second, instability frequency. This is the percentage of time neural activity and electric fields are simultaneously unstable. Both measures quantify the relative instability between fields and neural activity. Interestingly, we found that the values of both measures especially in the left hemisphere seemed to reflect HDRS scores.

The above results based on LFP data from the left hemisphere are in accord with converging evidence suggesting that the left hemisphere, shows consistent and specific dynamics in depression: early PET imaging research and post-stroke depression studies reveal that left relative to right frontal metabolic activity is a hallmark of depressive states^63,64^. Subsequent neuroimaging and lesion studies demonstrate that damage to or hypoactivity in the left DLPFC correlates strongly with the severity of depression symptoms, underscoring its crucial role in mood regulation^65^. Interventions such as repetitive transcranial magnetic stimulation (TMS) frequently target the left DLPFC, especially with a focus on its connectivity to SCC, which has emerged as target circuit for therapeutic modulation^66^. Acute local field potential (LFP) measurements show robust bilateral changes but reveal that only left beta power correlates with clinical depression scales (Hamilton) in the week after stimulation ^2^. Surveys of behavioral outcomes with bilateral stimulation consistently find stronger and more reliable responses with left over right hemisphere targeting^67^. New high-density intracranial cohorts continue to show that the left beta frequency serves as the most reliable biomarker of depressive state recovery, while the right hemisphere remains variable and may underlie aspects of affective or clinical instability^1,68^.

In previous work ^4,30^, one of us considered ephaptic field effects on memory networks. This revealed the importance of top down electric field effects similar to the ones observed here. In ^11^ electric fields (EFs) generated by FEF neural ensembles described by deep neural fields^69^ were found to exhibit greater stability than neural activity. EFs also had a relatively richer informational content. That work suggested that stability allows the brain to compute latent variables underlying the maintenance of the same memory in different trials. In ^30^, it was shown that stability follows from the theory of Complex Systems and that ephaptic coupling is a special case of a general theoretical result known as the *slaving principle*. Further, it was shown that the electric field is a *control* parameter: it evolves more slowly and constrains order parameters (like connectivity components ^69^) and enslaved parts (like spiking neurons ^57,70^). The slaving principle, borrowed from physics and biology, suggests that varying a control parameter, such as the electric field, induces various spatial patterns of neural responses, like those observed in cognitive maps ^71^. It also suggests a separation of timescales, where the electric field evolves more slowly than neural activity –which agrees with a common assumption in bio-electromagnetism, that the electric field is assumed to be quasi-static and electromagnetic propagation effects can be ignored ^31^. Similarly, correlations of single trial estimates of electric fields were found to be higher than correlations of similar neural activity estimates ^11^. Also, Granger causality (GC) strengths measuring top down influence from electric fields to neural activity were larger and more stable (varied much less) than strengths characterising bottom up influence from neural activity to field^30^.

Focusing on top down organizing influences (and the lack thereof), where electric fields affect neural activity to achieve efficient information processing is a fundamental shift from the classical neural coding principle^72^, where information is encoded and transmitted information via spike patterns. The classical coding principle suggests spike patterns allow the brain to make sense of complex sensory inputs, form memories, make decisions, and generate behavior, forming the foundation for intelligent action and thought. The cytoelectric coupling hypothesis^4^ on the other hand suggests that while for processing information spikes are crucial, higher level cognition requires neuron coordination through the collective influence of the brain’s aggregate electric field. Also, specific spatial-temporal electric field patterns have been associated with consciousness itself ^73–77^. Electric fields are thought to implement conscious awareness ^78,79^. At the same time depression can be thought of as a distinct conscious state with its own sensory content, including aberrant interoception and bodily experiences (e.g., feelings of fatigue, tiredness, lethargy ^80^), cognitive content, such as impairments in executive control and attention (e.g., difficulties with focus or decision-making), and temporal perception, encompassing an altered sense of presence^78^, quanta of time ^81^, and the perception of others ^82^. These ideas are consistent with the cytoelectric coupling hypothesis, which proposes that the re-emergence of depressive symptoms may reflect disruptions in electric field dynamics. From this perspective, the efficacy of DBS in depression may depend on its capacity to restore or stabilize electric fields, thereby reinstating coherent information processing and normal conscious function.

All in all, we used complex systems theory to shed light on brain power fluctuations in depression patients. Our interest in brain power was motivated by the fact that it seems to predict HDRS scores^1^. Power fluctuations seemed to be an epiphenomenon of itinerancy, that is, brain activity while neural circuits move between different energy states. We also suggested that power fluctuation itinerancy might be characterized using Lyapunov exponent measures (relative wandering time, instability frequency) that seemed to reflect HDRS scores. Whether the new measures reflect general mechanisms of rapid antidepressant action remains to be tested.

## Acknowledgements

This work was supported by the National Institutes of Health (ClinicalTrials.gov ID NCT01984710), the BRAIN Initiative through the National Institute of Neurological Disorders and Stroke (UH3NS103550,UH3NS141080).DAP was supported by the UK Medical Research Council (MRC) (Grant Number MR/W011751/1). SA was supported in part by the National Center for Advancing Translational Sciences of the National Institutes of Health under Award Number UL1TR002378 and KL2TR00238. PS was supported by the National Center for Advancing Translational Sciences of the National Institutes of Health under Award Number UL1TR002378 and TL1TR002382. The content is solely the responsibility of the authors and does not necessarily represent the official views of the National Institutes of Health.

---

**Supplementary Figure 1.**
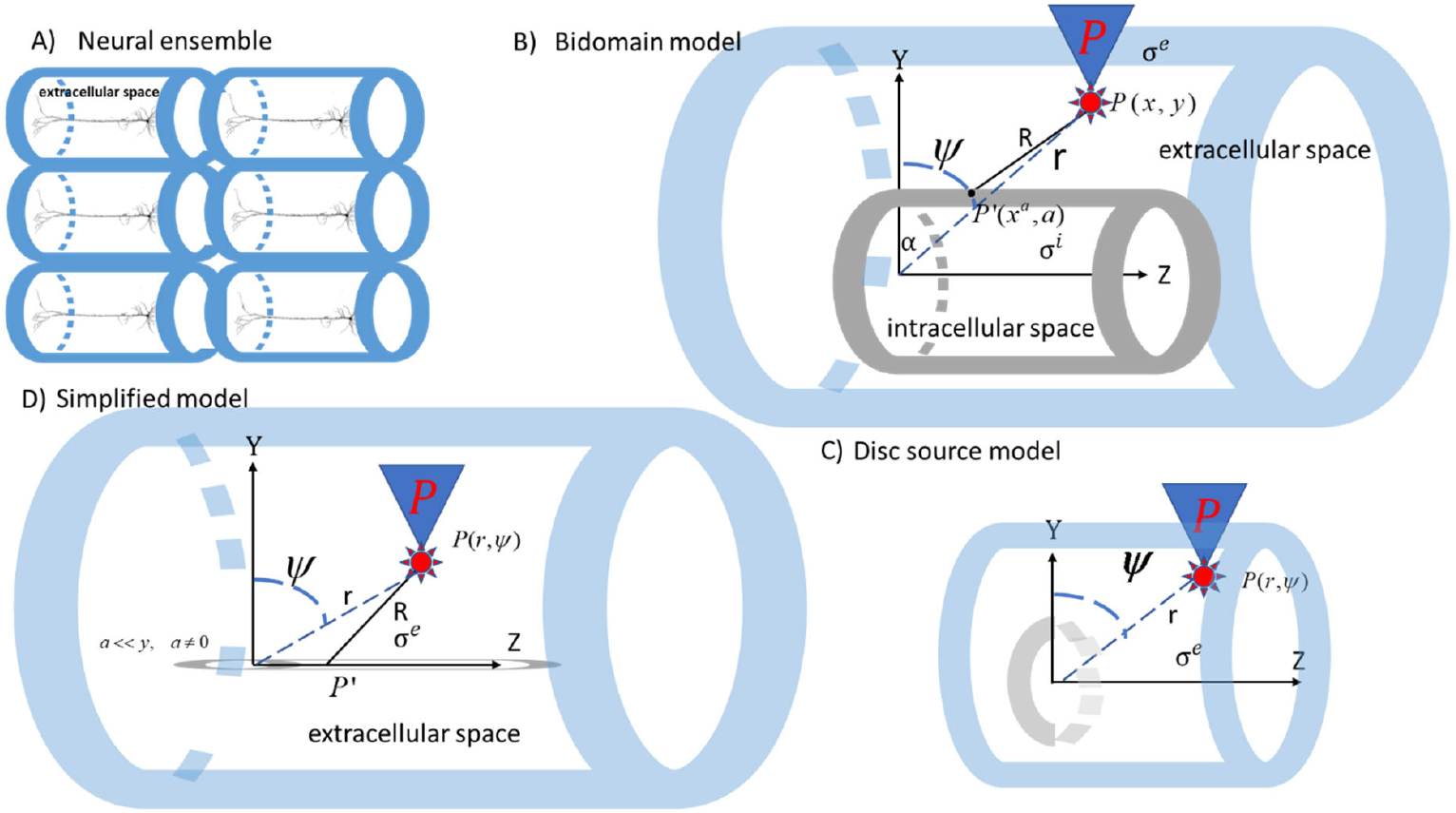

## Supplementary: Bidomain model of the ensemble electric field

We here summarize the bidomain model discussed in our earlier work ^11^. According to the theory of electromagnetism, the discontinuity between extracellular and intracellular potential gives rise to dipole sources with moments ^29^

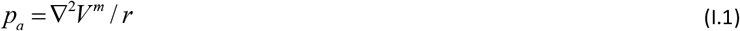

Here *p*_*a*_ is the moment of a neural ensemble whose center is at location (*x*_*a*_, *y*_*a*_), *r* is the brain resistivity with *r* = 2.2 Ohm ^83^ and we have assumed that the number of neurons is large and that each cell is very small compared to the distance at which the LFP electrode is placed. Also, the current density *I* ^*a*^ (*x*_*a*_, *y*_*a*_) that results from EPSPs and IPSPs is given by

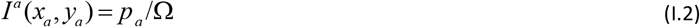

where Ω is the total volume of the ensemble. Neglecting ephaptic interactions *V* ^*m*^ ≈ *V* ^*e*^, and the extracellular electric potential generated by the current density *I* ^*a*^ (*x*_*a*_, *y*_*a*_) is given by

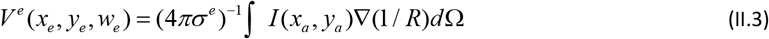

where *σ* ^*e*^ is the conductivity of the extracellular space, and *R* is the distance between the current source at the point *P* ‘(*x*_*a*_, *a*) of the neural ensemble and the point (*x, y*) in the extracellular space where we measure *V* ^*e*^, i.e. the location of the LFP electrode, 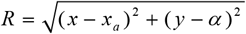 see Supplementary Figure 1B. Here, *a* is the radius of the grey cylinder in the figure. According to the bidomain model, Equation (I.3) can be written as ^84,85^

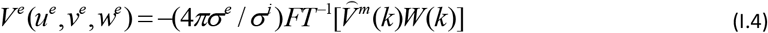

Where 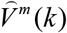 is the Fourier Transform of the transmembrane potential *V* ^*m*^ and *FT*^-*1*^ is its inverse Fourier Transform, that is,

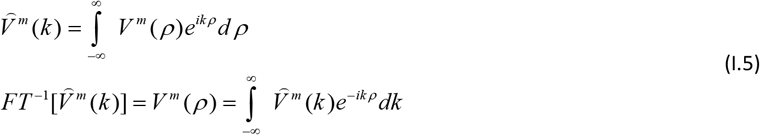

Equation (I.4) can then be written as

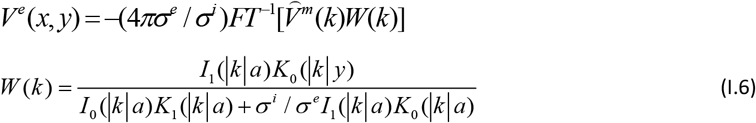

Where, *σ* ^*l*^, *l* ={*e,i*}, are the extra-and intra-cellular space conductivities and *I*_0_ (*y*), *I*_1_ (*y*), *K*_0_(*y*), *K*_1_(*y*) are modified Bessel functions of the first and second kind ^86^. Equation (I.6) is known as the bidomain model of the electric field.

